# Early events of the endophytic symbiotic between *Oryza sativa* and *Nostoc punctiforme* involve the SYM pathway

**DOI:** 10.1101/2021.04.09.436957

**Authors:** Consolación Álvarez, Manuel Brenes-Álvarez, Fernando P. Molina-Heredia, Vicente Mariscal

## Abstract

Symbiosis between cyanobacteria and plants is considered pivotal for biological nitrogen deposition in terrestrial ecosystems. Despite the large knowledge in the ecology of plant-cyanobacteria symbioses, little is known about the molecular mechanisms involved in the crosstalk between partners. A SWATH-mass spectrometry has been used to analyse, at the same time, the differential proteome of *Oryza sativa* and *Nostoc punctiforme* during the first events of the symbiosis. *N. punctiforme* activates the expression of thousands of proteins involved in signal transduction and cell wall remodelling, as well as 11 Nod-like proteins that might be involved in the synthesis of cyanobacterial-specific Nod factors. In *O. sativa* the differential protein expression was connected to a plethora of biological functions including signal transduction, defense-related proteins, biosynthesis of flavonoids and cell wall modification. *N. punctiforme* symbiotic inspection of *O. sativa* mutants in the SYM pathway reveals the involvement of this ancestral symbiotic pathway in the symbiosis between the cyanobacterium and the plant.

## 1. INTRODUCTION

*Nostoc punctiforme* (hereafter *Nostoc*) is one of the most versatile N_2_-fixing cyanobacteria. It occurs as free-living forms or in symbioses with plants from the four major phylogenetic divisions of terrestrial plants, reflecting a high diversity and low host specificity in its symbiotic interactions (Svenning et al., 2005; Adams et al., 2013; Warsham et al., 2018). *Nostoc* provides different modes of symbiotic associations with their host plants. It grows epiphytically in specialized compartments of liverworts, hornworts and *Azolla*, meanwhile colonizes endophytically stem glands of *Gunnera* sp. *Oryza sativa* (hereafter *Oryza*) and *Triticum vulgare*, providing fixed nitrogen to the plant (Gantar et al., 1991; Santi et al. 2013; Álvarez et al., 2020). Symbiotic nitrogen fixation with cereals is central for exploring new sustainable agricultural practices that may reduce the usage of synthetic fertilizers, whose application results in adverse environmental consequences (diCenzo et al., 2020). However, despite the beneficial effects of cyanobacterial nitrogen fixation in terrestrial ecosystems, to date little is known about the signalling mechanisms, crosstalk between partners and metabolic adaptations underlying the symbiotic process.

Independently of the mode of association, symbiotic interaction of *Nostoc* with plants comprise an early phase of interaction, which includes chemical signalling between partners, followed by a later phase where the cyanobacteria are physically associated with the host and supply nutrients to the plant. In response to nitrogen limitation, the plant produces signals that trigger hormogonia differentiation (which are the cyanobacterial infection units) and others compounds that act as chemoattractants (Meeks and Elhay, 2002; Nilson et al., 2006). Most of the knowledge on colonization steps and maintenance of *Nostoc*–plant symbioses are based on the *Nostoc* response to the liverwort *Blasia*, the hornworts *Anthoceros* and *Phaeoceros*, and the angiosperm *Gunnera* (Adams et al., 2013). Key cyanobacterial genes required for symbiosis have been predicted *in silico* (Warshan et al., 2017; 2018) and experimentally identified by proteomic and transcriptomic analyses (Ekman et al., 2006; Campbell et al., 2008; Wharshan et al., 2017). They include genes encoding proteins involved in chemotaxis and motility, oxidative stress response, transport of phosphate, amino acids and ammonium, and repression of photosynthetic CO_2_ fixation. These changes imply that the cyanobacterium shifts from photo-autotrophic to heterotrophic lifestyle, relying of the carbon provided by the host to sustain N_2_-fixation (Meeks and Elhai, 2002). Very little is known about the plant changes in response to the cyanobacterium. Expression changes in response to *Nostoc* have been studied by RNA-seq in *Anthoceros* and *Arabidopsis* (Li et al., 2020; Belton et al., 2020). Activation of receptor kinases and genes involved in stress response have been reported.

In other plant-microbial endosymbiosis, including the legume-*Rhizobium*, *Parasponia*-*Rhizobium*, actinorhizal plants-*Frankia* and arbuscular mycorrhizal (AM) fungi (Glomeromycota) symbioses, molecular genetic studies have revealed that signalling pathways of host plants largely overlap (Geurts et al., 2016; Oldroyd, 2013; Radhakrishnan et al., 2020). These signalling pathways comprise a well conserved network in land plants known as the ‘common symbiosis signalling pathway’ (SYM) (Oldroyd, 2013). The SYM pathway encompasses plasma membrane-localized LysM-type and LRR-type receptor kinases, a transcriptional network involving two predicted cation channels, CASTOR and POLLUX, the calcium/calmodulin-dependent protein kinase CCAMK, and CYCLOPS, which is phosphorylated by CCAMK (Parniske, 2008). This pathway is present in *Oryza* and is essential to AM symbiosis (Gutjar et al., 2009). SYM pathway is activated by Nod and mycorrhizal (Myc) factors, produced by rhizobia or AM, respectively (Oldroyd, 2013). They are chitooligosaccharides containing a pentameric chitin backbone and specific acyl groups that give plant selectivity (Oldroyd, 2013). Neither Nod factors nor Myc factors have never been identified in a cyanobacterium. However, *Nostoc* DNA sequences homologous to the rhizobial *nod* genes were identified two decades ago by heterologous hybridization (Rasmussen et al., 1996).

In contrast to the extensive knowledge of the signalling mechanisms in *Rhizobium*-legume symbioses, none of the signalling networks involved in *Nostoc* symbioses have been identified. In the present work, we have evaluated changes that occur in *Oryza* and *Nostoc* at an early phase of recognition (1 day) and a later phase of contact (7 days), providing a valuable overview of the recognition process. These changes have been determined by sequential window acquisition of all theoretical mass spectra (SWATH-MS) (Huang et al., 2015; Liu et al., 2019). This method is able to do label-free quantification in an MRM-like manner, providing the expression profile of a protein at proteome scale. We provide, for the first time, simultaneous temporal regulation recognition pathways in both the plant and the cyanobacterium during co-culture. We have identified a total of 2906 proteins, 1397 from *Nostoc* and 1509 from *Oryza.* Analysis of differentially expressed proteins revealed metabolic changes involved in adaptation to symbiosis when both partners were in contact. Analysis of *Nostoc* colonization on different *Oryza* mutants in the SYM pathway reveals the involvement of this common symbiotic pathway in the symbiosis between *Nostoc* and *Oryza*.

## 2. METHODS

### 2.1. Organisms and growth conditions

Rice seedlings (*Oryza sativa* L.) var. Puntal (Indica) were used for the proteomic analysis and the colonization inspection. The Nipponbare background was used for lines mutated in POLLUX (lines NC6423 and ND5050, *pollux*-2 and *pollux*-3, respectively), in CCAMK (lines NE1115 for *ccamk*-1 and NF8513 for *ccamk*-2, respectively) and in CYCLOPS (lines NC2415, and NC2713 for *cyclops*-2, and *cyclops*-3, respectively) (Gutjahr, et al 2008). All of them were kindly provided by Prof. Uta Paszkowski, University of Cambridge.

Rice seedlings were surface-sterilized by washing first with distilled water and then with 0.5% (w/v) calcium hypochlorite for 20 min. Later, the seeds were thoroughly washed with sterilized tap water and kept for germination on a wet filter paper in a container. Rice plants were grown hydroponically in 50-ml conical tubes with BG11_0_ (-N) medium (free of combined nitrogen) (Rippka et al. 1979). Experiments were carried out in a growth chamber at 28 °C, 75% relative humidity, 12 h light-12 h dark cycles and light intensity of 50 μmol·m^−2^·s^−1^.

*Nostoc punctiforme* PCC 73102 (also known as ATCC 29133) was obtained from the Biological Cultures Service of the IBVF, and grown in BG11_0_ medium supplemented with 2.5 mM NH_4_Cl, 5 mM TES-NaOH buffer (pH 7.5) at 30 °C in continuous light (50 μmol·m^−2^·s^−1^) in shaken liquid cultures (100 rpm) or on medium solidified with 1% (w/v) Difco agar. In order to avoid hormogonia differentiation, filaments of *Nostoc* were grown to a concentration of 2-3 μg Chl·ml^−1^ in BG11_0_ medium supplemented with 2.5 mM NH_4_Cl, 5 mM TES-NaOH buffer (pH 7.5) and 4 mM sucralose (Splitt and Risser 2016). Induction of hormogonia differentiation was carried out by washing and incubated in BG11_0_ without sucralose for 16 h.

### 2.2. Co-cultivation of *Nostoc* and rice plants

Seedlings of rice grown for 7 days were suspended in 50-ml conical tubes with BG11_0_ medium. After 4–5 days of acclimation, *Nostoc* inoculants containing hormogonia were added to the solutions at a final concentration of 0.8 μg Chl·ml^−1^. Co-cultivation was carried out in a growth chamber up to 35 days as described above for rice plants. Cultures of each partners were also grown separately in parallel as controls.

To determine association to plant roots, Chlorophyll *a* content was measured as in Álvarez et al., 2020, according to Arnon (1949). Root samples were directly taken at different times of co-culture. Chlorophyll was referred to the roots fresh weight.

### 2.3. Sample preparation for confocal microscopy

Fresh rice roots were cut and washed intensively with tap water. Alternatively, root slices were obtained by excision with a blade. Samples were mounted in a coverslip and examined with a Leica TCS SP2 confocal microscope using HC PL-APO CS ×20 or HCX PLAM-APO ×63 1.4 NA oil immersion objectives. Cyanobacterial autofluorescence was excited with 488 nm light irradiation from an Argon laser, and fluorescence emission was monitored across windows of 650–700 nm. Root’s lignin and suberin autofluorescence was similarly excited with 476 and 488 nm laser irradiation, and fluorescence emission was monitored across windows of 510–533 nm. Z series containing from 90 to 130 frames were stacked and processed with the Image J program (version 1.41).

### 2.4. Protein extraction and trypsin digestion

Total protein extracts were prepared from biological triplicates at 1 day and 7 days post inoculation from control *Nostoc* samples (5 ml of culture at 50 μg Chl·ml^−1^ per replicates), control *Oryza* root samples (300 mg from 15 plants per replicate) and *Nostoc* treated plant roots (300 mg from 15 plants per replicate). Samples were ground to powder in liquid nitrogen and then incubated in lysis buffer (50 mM Tris-Cl pH 7.5, 1 mM PMSF (phenylmethanesulfonylfluoride), 1 mM EDTA (ethylenediaminetetraacetic acid), 2% SDS and Complete™ mini EDTA free protease inhibitor cocktail) and centrifuged at 16,000 ×*g* at 4 °C for 15 min. Protein concentrations were determined by Bradford method (Bio-Rad).

Protein samples were precipitated by TCA/acetone. Precipitated samples were resuspended in 50 mM ammonium bicarbonate with 0.2% Rapigest (Waters) for protein determination. Protein samples (20 μg) were digested with trypsin as described previously (Vowinckel et al., 2014).

### 2.5. Proteome analysis by SWATH-MS

The SWATH-MS analysis was performed at the Proteomic Facility of the Institute of Plant Biochemistry and Photosynthesis, Seville, Spain. A data-dependent acquisition (DDA) approach using nano-LC-MS/MS was initially performed to generate the SWATH-MS spectral libraries.

Peptide and protein identifications were performed using Protein Pilot software (version 5.0.1, Sciex) with the Paragon algorithm. The search was conducted against Uniprot *Oryza* proteome, Uniprot *Nostoc* proteome or a combined database with Uniprot *Oryza*+*Nostoc* proteome. Automatically generated reports in ProteinPilot were manually inspected for FDR (false discovery rate) cut-off proteins. Only proteins identified at FDR ≤1% were considered for output and subsequent analysis. Protein-specific peptide and peptides not shared between the two organisms were selected for further analysis. The combined *Oryza*+*Nostoc* library was used to detect interspecies peptides. Peptides shared by *Oryza* and *Nostoc* were not used in the downstream analysis. As a result of this process, we generated two spectral libraries, one for *Oryza* and one for *Nostoc* containing species-specific peptides.

For relative quantification using SWATH analysis, each biological replicate was quantified using three technical replicates and a data-independent acquisition (DIA) method. Each sample (1 μg of protein) was analysed using SWATH-MS acquisition method on the LC-MS equipment with a LC gradient. The method consisted of repeated acquisition cycles of TOF MS/MS scans (230 to 1500 m/z, 60 ms acquisition time) of 60 overlapping sequential precursor isolation windows of variable width (1 m/z overlap) covering the 400-1250 m/z mass range from a previous TOF MS scan (400-1250 m/z, 50 ms acquisition time) for each cycle. The total cycle time was 3.7 s. Autocalibration of the equipment and chromatographic conditions were controlled by an injection of a standard of digested β-galactosidase from *Escherichia coli* between the replicates.

SWATH MS spectra alignment was performed with the PeakView 2.2 (Sciex) software with the MicroApp SWATH 2.0 using the *Oryza* or *Nostoc* spectral libraries generated as described above.

After data processing, the processed mrkw files containing protein information from PeakView were loaded into MarkerView (Version 1.2.1.1, AB Sciex) for normalization of protein intensity (total area sums) using the built-in total ion intensity sum plug-in.

Mass spectrometry raw proteomic data have been deposited to the ProteomeXchange Consortium via the PRIDE partner repository with the identifier PXD022229.

### 2.5. Computational methods

Quality of normalised data was analysed using PCA analysis and results were visualized using *ggbiplot* R package. Normalised data were later processed with *limma* R package (Ritchie et al., 2015) to extract changes in protein abundance. We performed the following comparisons between the cyanobacterial samples: *Nostoc* plus *Oryza* after 24 h (*Nostoc_Oryza*_24h) versus *Nostoc* without plant after 24h (*Nostoc*_24h), *Nostoc* plus *Oryza* after 7 days (*Nostoc_Oryza* _7days) versus *Nostoc* without plant after at 7 days (*Nostoc*_7days), *Nostoc* plus *Oryza* after 7 days (*Nostoc_Oryza*_7days) versus *Nostoc* plus *Oryza* after 24 h (*Nostoc_Oryza*_24h) and *Nostoc* without *Oryza* after 7 days (*Nostoc*_7days) versus *Nostoc* without *Oryza* after 24 h (*Nostoc*_24h). The same logic comparisons were followed for the plant samples. A total of 1076 cyanobacterial proteins had a statistically differential abundance in at least one comparison (fold change > 1.5 or < 0.66667, and adjust p-value < 0.05). In the case of the plant, a total of 736 plant proteins fulfilled the same criteria.

Clustering analysis was carried out using *mfuzz* R package (Kumar and Futschik, 2007) using a ‘fuzzifier’ of 1.25647 (m = 1.25647) for *Nostoc* and 1.265078 (m = 1. 265078) for *Oryza*. Cyanobacterial or plant proteins considered to be in the core of a cluster were selected using a strict confidence threshold of 0.80.

## 3. RESULTS

### 3.1. Roots of *Oryza sativa* sp. Indica are endophytically colonized by *Nostoc punctiforme*

Association of *Nostoc* to *Oryza* roots was first studied. A fixed amount of *Nostoc* hormogonia containing 0.8 μg Chl·ml^−1^ was inoculated to hydroponic cultures of *Oryza* plants. Chlorophyll of the roots was measured at different time points ranging from 24 h to 35 days after inoculation (Figure 1A). *Nostoc* hormogonia reached the plant within few hours after inoculation, being strongly attached to the plant roots (Figure 1B and C). The cyanobacterium started growing in the proximity of the roots, increasing the biomass until 15 days. At 35 days post inoculation colonization of the plant was evident, being epidermal cells and radical hairs entirely colonized (Figure 1C). At this time, a significant difference was observed in length and fresh weight of plants treated with *Nostoc* (Figure S1).

**Figure 1.**
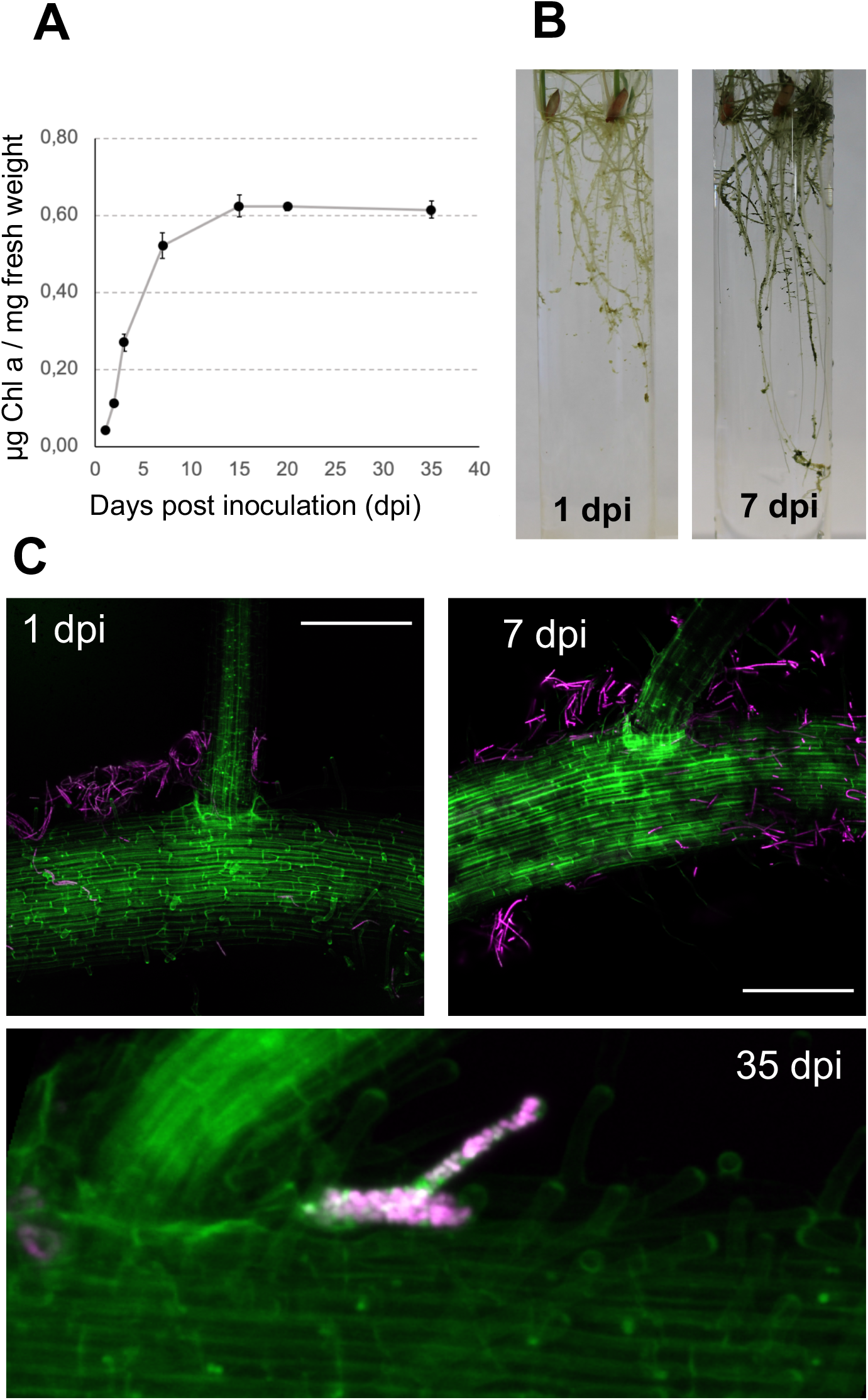
Association of *Nostoc* to *Oryza* sp. Indica roots. A) Cyanobacterial association to rice roots, quantified as μg chlorophyll *a* (Chl*a*)·mg^−1^ root weight. The values are the means ± SE. B) Appearance of rice roots inoculated with *Nostoc* at 1 and 7 days post inoculation (dpi). C) Rice roots inoculated with *Nostoc* at 1, 7, and 35 days post inoculation (dpi) visualized by a confocal microscope. Stacks generated from 90 to 130 frames are shown. In green, suberin autofluorescence from the plant cell walls; in magenta, cyanobacterial chlorophyll autofluorescence. Scale bar: 100 μm.

### 3.2. Simultaneous quantitative proteomics of *Nostoc* and *Oryza*

To assess the proteome coverage of the early steps of recognition of the two partners, 1 day and 7 days of co-culture were selected for the quantitative analysis. Freshly hormogonia from *Nostoc* and *Oryza* seedlings were co-cultured in BG11_0_. As controls, cultures of each partners were grown separately in parallel. Total protein extracts from controls and *Nostoc* treated plants (in triplicate) at 1 day and 7 days post inoculation were analysed by SWATH (Figure 2). Among 2906 total identified proteins, 1397 proteins were from *Nostoc* and 1509 proteins belong to *Oryza.*

**Figure 2.**
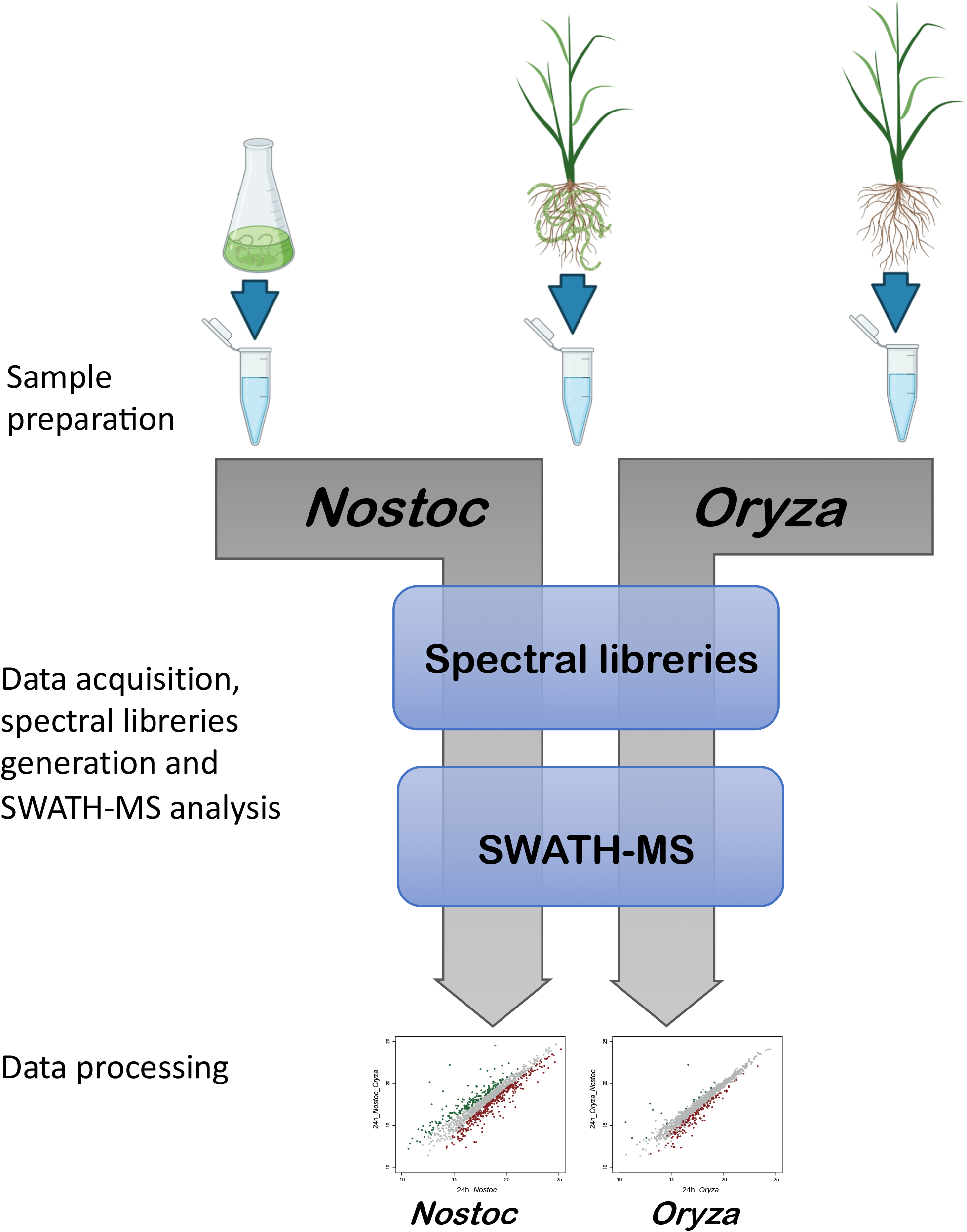
Overview the schematic workflow used for quantitative proteomic in *Nostoc* and *Oryza*. The workflow consists of a first step of sample preparation and data acquisition by LC-MS/MS for generation of species-specific spectral libraries. In a second step, relative quantification of protein changes was made by SWATH-MS analysis. The subsequent data normalization was made with *limma* R package to extract changes in protein abundance.

We performed pair-wise comparison to explore the similarities and differences in the corresponding proteomes (Supplementary File 1). As a result, 666 and 862 proteins were found to change their abundance at 1 day and 7 days post inoculation relative to non-inoculated samples, respectively (p-value ≤ 0.05), fold change ≥1.5 or < 0.66667). At 1 day, 159 plant proteins and 507 cyanobacterial proteins changed their abundance. After 7 days post-inoculation, 321 plant proteins and 541 cyanobacterial proteins had a differential accumulation (Figure 3). Among the 1397 proteins identified in *Nostoc,* 36% and 39% proteins changed their abundance at 1 day and 7 days after post-inoculation, respectively. However, only 10.5% and 21% of identified plant proteins had a differential accumulation after 1 day and 7 days of treatment. The higher number of differentially expressed proteins in *Nostoc* from the very first moment reveals profound changes in the active metabolism of the cyanobiont responding to the plant. In the case of the plant, most of the changes take place at 7 days after co-cultivation.

**Figure 3.**
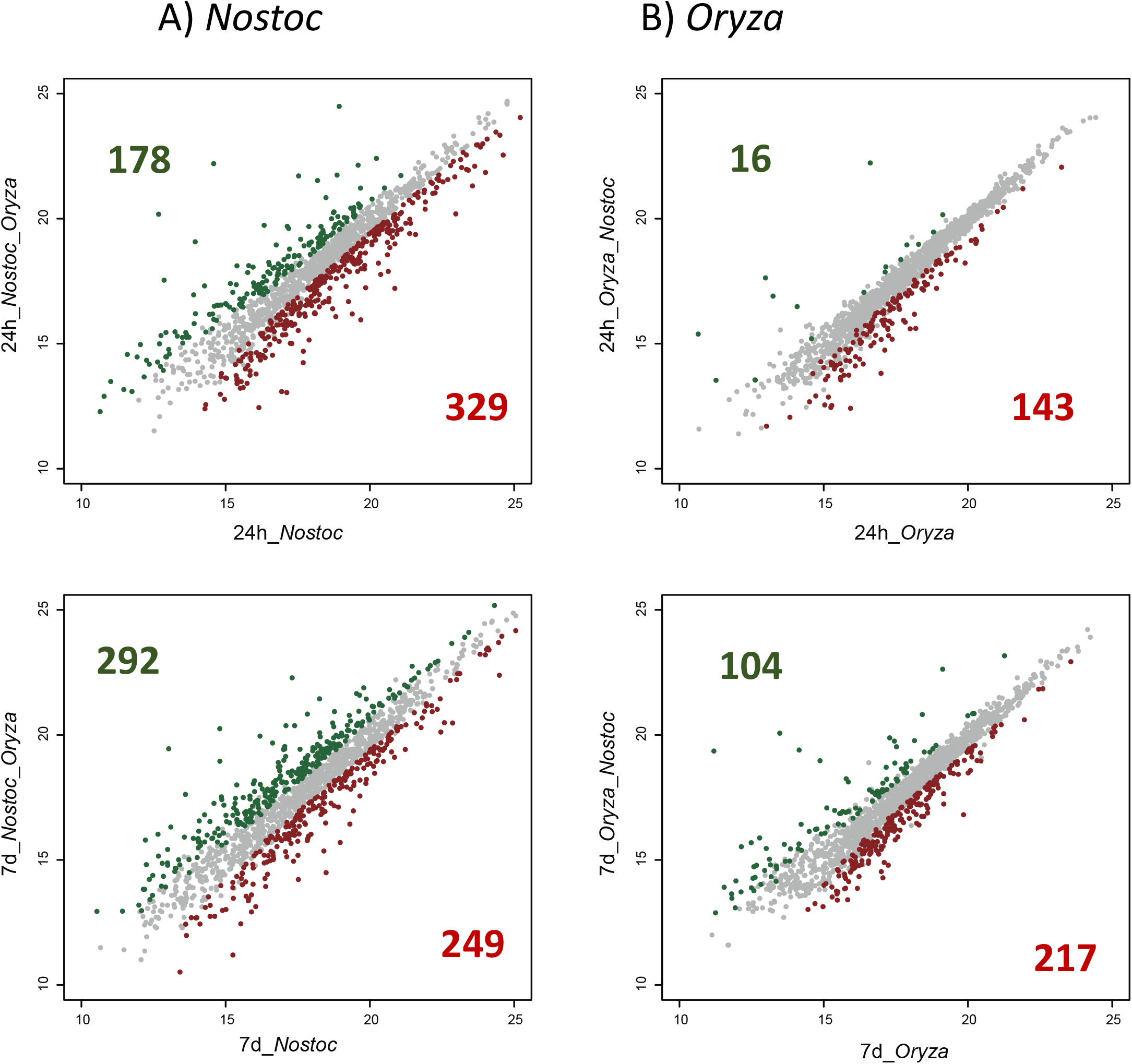
Global protein expression changes in *Oryza* and *Nostoc* during co-culture. Protein expression was analysed by *limma* R package. The number of differentially expressed proteins is shown in green, for proteins significantly upregulated and in red, for proteins significantly downregulated (fold change > 1.5 or < 0.66667, and adjust p-value < 0.05).

As an initial characterization and check of our experimental design, samples were analysed using principal component analysis (PCA). The PCA analysis showed a good quality data and no batch effect or confounding factors. Samples belonging to the different conditions and times segregated into well separated groups, and biological replicates clustered tightly together (Figure S2). No outlier values were detected, so all the replicates were used for further analyses.

In order to understand the timeline of the symbiosis as a whole, we performed pair-wise comparisons that consider not only the *Nostoc* treatment but also the time-course expression of proteins. In order to ascertain groups of proteins with the same regulation, a clustering analysis using *mfuzz* R package was carried out in both organisms (Kumar and Futschik, 2007). We obtained 8 clusters in *Nostoc* and 6 clusters in *Oryza* with at least 80% confidence (Figures S3 and S4). Proteins belonging to each cluster can be found at glance at the data files Supplementary File 1. A heat map representation of the proteins in each cluster illustrates the consistency of the clustering and between individual replicates of the treatments (Figure 4).

**Figure 4.**
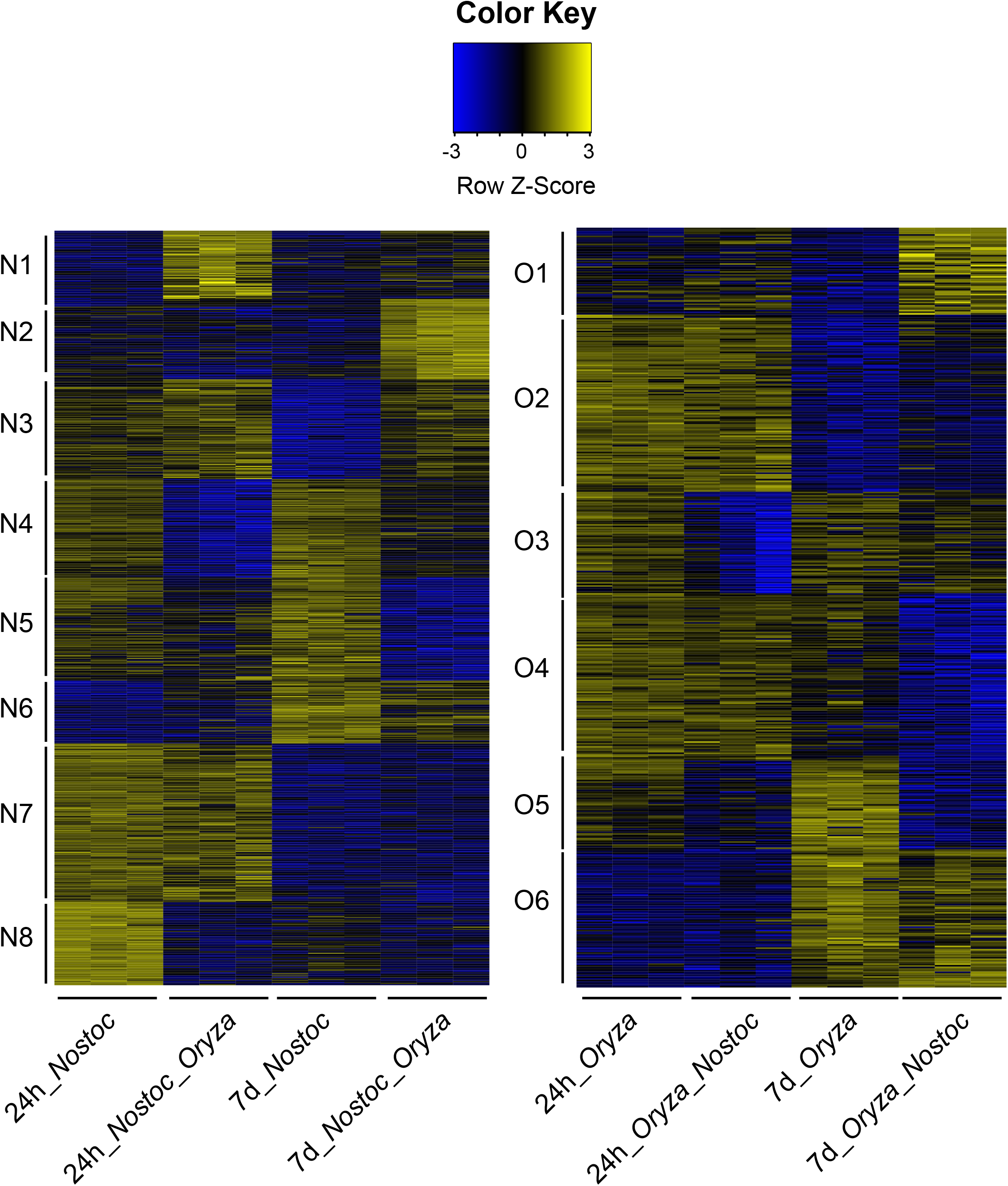
Heat map profiles of proteins differentially expressed in *Nostoc* and *Oryza*. Differentially expressed proteins were clustered into eight groups in *Nostoc* (N1 to N8), and six groups in *Oryza* (O1 to O6) based on their expression profile and *mfuzz* analysis. The legend shows Z-scores. The values were mean centred, and the colours scaled from −3 to +3 standard deviations.

### 3.3. Proteins up-regulated in *Nostoc* and *Oryza* in response to the partner

In *Nostoc*, 178 proteins were significantly induced at 1 day of co-culture (included in cluster N1) and 292 proteins were induced at 7 days of co-culture (included in clusters N2 and N3). From them, 81 proteins were induced both a 1 and 7 days in response to the plant partner. In *Oryza*, a total amount of 104 proteins were significantly induced in the presence of *Nostoc* (included in cluster O1 and O2). Most of the changes in the plant were observed at 7 days post inoculation, being only 16 proteins induced at 24 h. Proteins activated in both organisms can be found at glance at the data files Supplementary File 1 filtering by 1. A group of selected proteins is shown in Table 1.

**Table 1.** Selection of proteins significantly activated in *Oryza* and *Nostoc* at 1 day and 7 days of co-culture

| Organism | Anotation | Locus | FC 1d | P value | FC 7d | P value |
| --- | --- | --- | --- | --- | --- | --- |
| **Signaling** |  |  |  |  |  |  |
| ***Nostoc*** | PtxB | Npun_F2162 | **25.51** | 1.3E-06 | **3.47** | 3.4E-02 |
|  | PtxD | Npun_F2164 | 1.26 | 2.8E-01 | **1.83** | 4.5E-03 |
|  | PtxG | Npun_F2167 | 1.68 | 1.4E-01 | **2.30** | 1.8E-02 |
|  | HmpB | Npun_F5961 | 0.47 | 2.4E-04 | **2.24** | 7.9E-05 |
|  | GAF sensor | Npun_R1597 | **18.11** | 1.7E-07 | **3.91** | 4.2E-03 |
|  | GAF sensor | Npun_F2781 | 1.13 | 6.9E-01 | **2.32** | 2.0E-03 |
|  | Response regulator receiver | Npun_F0021 | **4.54** | 5.5E-05 | 0.88 | 7.6E-01 |
|  | Response regulator receiver | Npun_R0589 | **2.56** | 6.0E-04 | **2.16** | 3.9E-03 |
|  | Response regulator receiver | Npun_F6378 | 0.99 | 9.4E-01 | **2.50** | 9.5E-08 |
|  | Response regulator receiver | Npun_AR130 | 1.10 | 8.1E-01 | **2.34** | 1.3E-02 |
|  | cGMP-specific 3',5'-cyclic phosphodiesterase | Npun_R2479 | **5.15** | 1.9E-24 | **1.68** | 1.1E-10 |
| ***Oryza*** | E3 ubiquitin ligase | B8BET8 | 0.66 | 2.6E-01 | **3.93** | 4.5E-05 |
|  | SH3 domain-containing protein | B8AKM5 | 0.70 | 8.9E-02 | **2.17** | 1.4E-04 |
| | Importin-α | A2X9V6 | **5.29** | 1.1E-05 | **17.08** | 1.9E-10 |
| **Symbiosis-related proteins** |  |  |  |  |  |  |
| ***Nostoc*** | Putative NodB | Npun_F3876 | 1.22 | 7.6E-01 | **5.76** | 2.8E-03 |
|  | Putative NodE | Npun_R0041 | **8.34** | 6.3E-08 | 0.49 | 2.7E-02 |
|  | Putative NodE | Npun_R6590 | **2.13** | 1.8E-04 | **2.75** | 2.1E-06 |
|  | Putative NodE | Npun_F3359 | 1.05 | 6.3E-01 | **2.72** | 2.0E-13 |
|  | Putative NodE | Npun_F3360 | **1.85** | 6.9E-04 | **1.98** | 2.0E-04 |
|  | Putative NodO | Npun_R6244 | **2.53** | 4.6E-02 | **5.39** | 4.5E-04 |
|  | Putative NodO | Npun_F1217 | 0.71 | 1.2E-03 | **1.91** | 9.1E-08 |
|  | Putative NodO | Npun_DR001 | **3.98** | 4.9E-06 | 1.25 | 4.2E-01 |
|  | Putative NodO | Npun_R3354 | **1.65** | 3.9E-04 | 0.70 | 7.2E-03 |
|  | Putative NodO | Npun_F3370 | **1.78** | 2.1E-06 | 0.56 | 1.9E-06 |
|  | Putative NodO | Npun_F4679 | **1.57** | 3.7E-03 | 0.57 | 4.0E-04 |
| **Oryza** | Flavonoid 3'-monooxygenase | A2Z5X0 | 0.78 | 6.4E-01 | **2.23** | 3.6E-02 |
|  | Ankyrin_REP_REGION domain-containing protein | A2XP52 | 1.06 | 9.2E-01 | **2.20** | 8.5E-03 |
| **Cell adhesion** |  |  |  |  |  |  |
| **Nostoc** | FG-GAP | Npun_R5402 | **3.03** | 1.3E-03 | 1.08 | 8.6E-01 |
|  | FG-GAP | Npun_R1349 | **2.51** | 1.7E-06 | 0.61 | 4.0E-03 |
| **Oryza** | FL-AGP | A2YWR4 | **2.05** | 2.5E-01 | **3.10** | 3.6E-02 |
|  | FL-AGP | A2XUI8 | 0.78 | 4.8E-02 | **2.00** | 1.7E-07 |
|  | FL-AGP | A2YTW2 | 0.89 | 5.6E-01 | **1.64** | 1.8E-03 |
| **Cell wall modification** |  |  |  |  |  |  |
| **Nostoc** | Pullulanase | Npun_R0164 | 1.18 | 5.4E-02 | **2.23** | 1.5E-11 |
|  | β-glucosidase | Npun_F2285 | 0.8 | 4.4E-03 | **1.88** | 2.2E-10 |
|  | Pectate lyase | Npun_R5506 | 1.31 | 4.6E-04 | **1.52** | 8.0E-07 |
|  | N-acetylmuramoyl-L-alanine amidases | Npun_F1845 | **2.07** | 7.4E-07 | 1.16 | 2.5E-01 |
|  | N-acetylmuramoyl-L-alanine amidases | Npun_F6436 | 1.14 | 9.1E-03 | **1.82** | 4.3E-14 |
|  | N-acetylmuramoyl-L-alanine amidases | Npun_R6032 | **2.85** | 8.8E-03 | 1.48 | 3.3E-01 |
|  | N-acetylmuramidases | Npun_R4588 | **3.26** | 1.6E-06 | **17.72** | 3.0E-15 |
|  | N-acetylmuramidases | Npun_F5932 | 1.49 | 1.9E-06 | **2.09** | 4.5E-12 |
|  | N-acetylmuramic acid 6-phosphate hydrolase | Npun_R6016 | 0.74 | 9.4E-03 | **2.08** | 6.8E-08 |
|  | L-alanyl-D-glutamate peptidase | Npun_R4589 | 1.56 | 2.9E-01 | **3.64** | 1.7E-03 |
|  | Penicillin amidase | Npun_R3418 | **2.9** | 4.8E-03 | 1.04 | 9.4E-01 |
| **Oryza** | Cellulose synthase | A2YRG4 | 1.13 | 8.2E-01 | **2.28** | 1.4E-02 |
|  | Omega-hydroxypalmitate O-feruloyl transferase | A2YA31 | 1.16 | 7.6E-01 | **2.06** | 2.4E-02 |
|  | Putative endoglucanase | A2XV47 | 1.12 | 7.8E-01 | **2.2** | 3.7E-03 |
| **Defense response and oxidative stress** |  |  |  |  |  |  |
| **Nostoc** | OCP N-terminal domain-containing proteins | Npun_F6242 | **2.23** | 5.1E-07 | **13.55** | 2.7E-19 |
|  | OCP N-terminal domain-containing proteins | Npun_F6243 | 0.63 | 3.6E-01 | **10.36** | 9.3E-06 |
|  | OCP N-terminal domain-containing proteins | Npun_R5130 | 1.39 | 6.5E-02 | **5.33** | 1.7E-11 |
|  | OCP N-terminal domain-containing proteins | Npun_F5913 | 1.52 | 1.7E-01 | **4.34** | 9.2E-06 |
|  | Thioredoxin | Npun_R3962 | 0.95 | 6.5E-01 | **2.37** | 1.2E-09 |
|  | Thioredoxin | Npun_R5376 | 1.00 | 9.7E-01 | **1.70** | 1.3E-10 |
|  | Thioredoxin reductase | Npun_R0672 | 0.95 | 8.3E-01 | **1.59** | 2.6E-02 |
|  | Catalase | Npun_F0233 | 0.48 | 2.5E-02 | **3.19** | 5.9E-04 |
|  | Catalase | Npun_R4582 | 1.09 | 6.2E-01 | **2.60** | 1.5E-07 |
|  | Catalase | Npun_F5237 | 1.38 | 7.3E-07 | **2.12** | 3.9E-15 |
|  | Peroxidase | Npun_R5469 | **2.06** | 4.4E-06 | **3.01** | 1.0E-09 |
|  | Superoxide dismutase | Npun_F1605 | 0.95 | 6.8E-01 | **1.62** | 2.1E-05 |
| **Oryza** | Class III Peroxidases | A2Y667 | **1.58** | 1.9E-03 | 1.35 | 2.9E-02 |
|  | Class III Peroxidases | A2YHC0 | 0.84 | 8.5E-01 | **5.46** | 5.5E-03 |
|  | Class III Peroxidases | A2WZD6 | 0.99 | 9.4E-01 | **1.53** | 2.6E-04 |
|  | Glutathione S-transferases | A2Z9J9 | 0.87 | 9.0E-01 | **4.44** | 2.0E-02 |
|  | Glutathione S-transferases | Q6WSC3 | 0.86 | 6.2E-01 | **1.78** | 8.9E-03 |
|  | Glutathione S-transferases | B8A962 | **1.58** | 2.6E-03 | 0.67 | 5.6E-03 |
|  | Catalase | B8AGH7 | 1.08 | 4.8E-02 | **1.61** | 7.3E-15 |
|  | Gnk-2-homologous domain | A2XZM8 | 0.73 | 5.4E-01 | **3.75** | 8.9E-04 |

#### 3.4.1. Signalling

In *Nostoc*, we found a number of proteins highly induced in response to the plant. Some of them had been previously identified in other reports studying *Nostoc*-plant symbioses (Campbel et al., 2008; Warsham et al., 2017; Ekman et al., 2006). For example, we found three proteins (PtxB, PtxD and PtxG) of a system controlling positive phototaxis (Ptx system; Campbell et al., 2015), the response regulator receiver HmpB and two GAF sensor signal transduction histidine kinases (Npun_R1597 and Npun_F2781). Additionally, we found other proteins not described previously, including four response regulator receiver proteins (Npun_F0021, Npun_R0589, Npun_F6378 and Npun_AR130) and a putative cGMP-specific 3’,5’-cyclic phosphodiesterase (Npun_R2479). These sensor proteins might be involved in transducing plant signals involved in plant recognition. In *Oryza*, we found a predicted E3 ubiquitin ligase (B8BET8), a SH3 domain-containing protein (B8AKM5), two response regulators (B8AYY5 and B8AG68) and an Importin-α (A2X9V6).

#### 3.4.2. Cell adhesion and cell wall modification

*Nostoc* showed a strong adherence to the *Oryza* roots only a few hours after co-cultivation (Fig. 1). In accordance to this, we found upregulated *Nostoc* proteins involved in cell adhesion, including two FG-GAP repeat proteins (Npun_R5402 and Npun_R1349). FG-GAP proteins are involved in chemical and physical contact in the *Nostoc*-moss symbiosis (Warsham et al., 2017). In *Oryza*, three fasciclin-like arabinogalactan proteins (FLAs), a subclass of arabinogalactan proteins (AGPs), were significantly activated (A2YWR4, A2XUI8 and A2YTW2). FLAs.

Cell walls are deeply involved in the molecular talk between partners during plant-microbe interactions. We found a number of proteins annotated in the category of plant cell-wall modification, significantly induced at 7 days post inoculation. In *Oryza*, a cellulose synthase (A2YRG4), an expansin-like EG45 domain-containing protein (A2ZA45), a putative endoglucanase (A2XV47) and an omega-hydroxypalmitate O-feruloyl transferase (A2YA31), involved in the synthesis of the suberin polymer, were significantly induced. In *Nostoc*, they included a pullulanase (1,4-alpha-glucan branching enzyme GlgB, Npun_R0164), a putative ß-glucosidase (Npun_F2285) and a pectate lyase (Npun_R5506). The later was also upregulated in the exoproteome of *Nostoc* in contact with *Pleurozium schreberi* (Warsham et al., 2017). The activation of these enzymes might reveal a remodelling of the cell plant wall in the presence of the host, as already reported in the *Medicago*-AM symbiosis (Siciliano et al., 2007).

Induction of proteins involved in bacterial peptidoglycan recycling and modulation was also detected in *Nostoc*. We found three N-acetylmuramoyl-L-alanine amidases (AmiC1, Npun_F6436 and Npun_R6032), two N-acetylmuramidases (Npun_R4588 and Npun_F5932), a N-acetylmuramic acid 6-phosphate hydrolase (Npun_R6016), a L-alanyl-D-glutamate peptidase (Npun_R4589) and a penicillin amidase (Npun_R3418). Up-regulation of these peptidoglycan-modifying enzymes is indicative of strong physiological changes in the cyanobacterium that might be related to the adaptation to the symbiotic lifestyle.

#### 3.4.3. Defense response and oxidative stress

Convergence between signalling mechanisms in plant pathogenesis and symbiosis is known. Comparison of the two interactions has revealed that plants appear to detect both pathogenic and symbiotic microbes by a similar set of genes (Zhao et al., 2008). We detected in *Oryza* a number of significantly activated proteins involved in defense response. They included three class III peroxidases (A2Y667, A2YHC0 and A2WZD6), three glutathione S-transferases (A2Z9J9, Q6WSC3 and B8A962), a catalase (B8AGH7) and a Gnk2-homologous domain-containing protein (A2XZM8).

In *Nostoc*, a number of proteins involved in response to oxidative stress were significantly induced mostly at 7 days in co-culture with the plant. They included two thioredoxins (Npun_R3962 and Npun_R5376), a thioredoxin reductase (Npun_R0672), three catalases (Npun_F0233, Npun_R4582 and Npun_F5237), a peroxidase (Npun_R5469), a superoxide dismutase (Npun_F1605), a flavorubredoxin (Npun_F4866) and four N-terminal domain (NTD) of the photoactive orange carotenoid (OCP)-like proteins (NTD-like proteins) (Npun_F6242, Npun_F6243, Npun_R5130 and Npun_F5913). NTD-like proteins are involved in photoprotection and response to oxidative stress. It is well known that catalases are essential for the establishment of plant-microbial symbioses (Jamet et al., 2003). In that sense, it is important to note that Npun_R4582 and Npun_F5237 were also activated in the cyanobacterium in symbiosis with bryophytes (Ekman et al., 2006; Warshan et al., 2017).

#### 3.4.4. Plant flavonoids and cyanobacterial Nod factors

Flavonoids are crucial signalling molecules in the symbiosis between legumes and their nitrogen-fixing symbionts (Liu et al, 2016). In *Oryza*, we found a putative flavonoid 3’-monooxygenase (A2Z5X0) involved in the eriodyctiol biosynthesis from naringenin. Naringenin is exuded by some legume roots and also acts as an inducer of *nod* genes in rhizobia (Novak et al., 2002). We have found significantly activated in our proteomic analysis a number of *Nostoc* proteins with homology to Nod factor biosynthetic enzymes from *Rhizobium* and *Frankia.* They included a putative NodB polysaccharide deacetylase (Npun_F3876), four putative NodE beta-ketoacyl synthases (Npun_R0041, Npun_R6590, Npun_F3359 and Npun_F3360) and six proteins with homology to NodO (Npun_R6244, Npun_F1217, Npun_DR001, Npun_R3354, Npun_F3370 and Npun_F4679).

### 3.5. The *Oryza* SYM pathway is required for *Nostoc* colonization

Nod factors (in *Rhizobium*) and MyC factors (in AM) induce symbiotic responses specifically on roots of the plant hosts through the common symbiosis (SYM) pathway (Streng et al, 2011). In *Oryza*, components of the SYM pathway have been already identified (Gutjahr et al., 2008). In order to know the involvement of the SYM pathway in the *Nostoc-Oryza* symbiosis, homozygous *Oryza* mutant lines with insertions in SYM signalling components upstream and downstream of Ca^2+^ spiking, were analysed in *Nostoc* colonization. They included two predicted cation channels, CASTOR and POLLUX, the calcium/calmodulin-dependent protein kinase CCAMK, and CYCLOPS which is phosphorylated by CCAMK. Thus, *Nostoc* hormogonia were inoculated to plants of the different mutants and colonization was evaluated by means of confocal microscopy. We found that colonization was severely affected in all the mutants tested (Figure 5). Their deficient symbiotic phenotypes are consistent with a crucial role of the common SYM pathway in rice.

**Figure 5.**
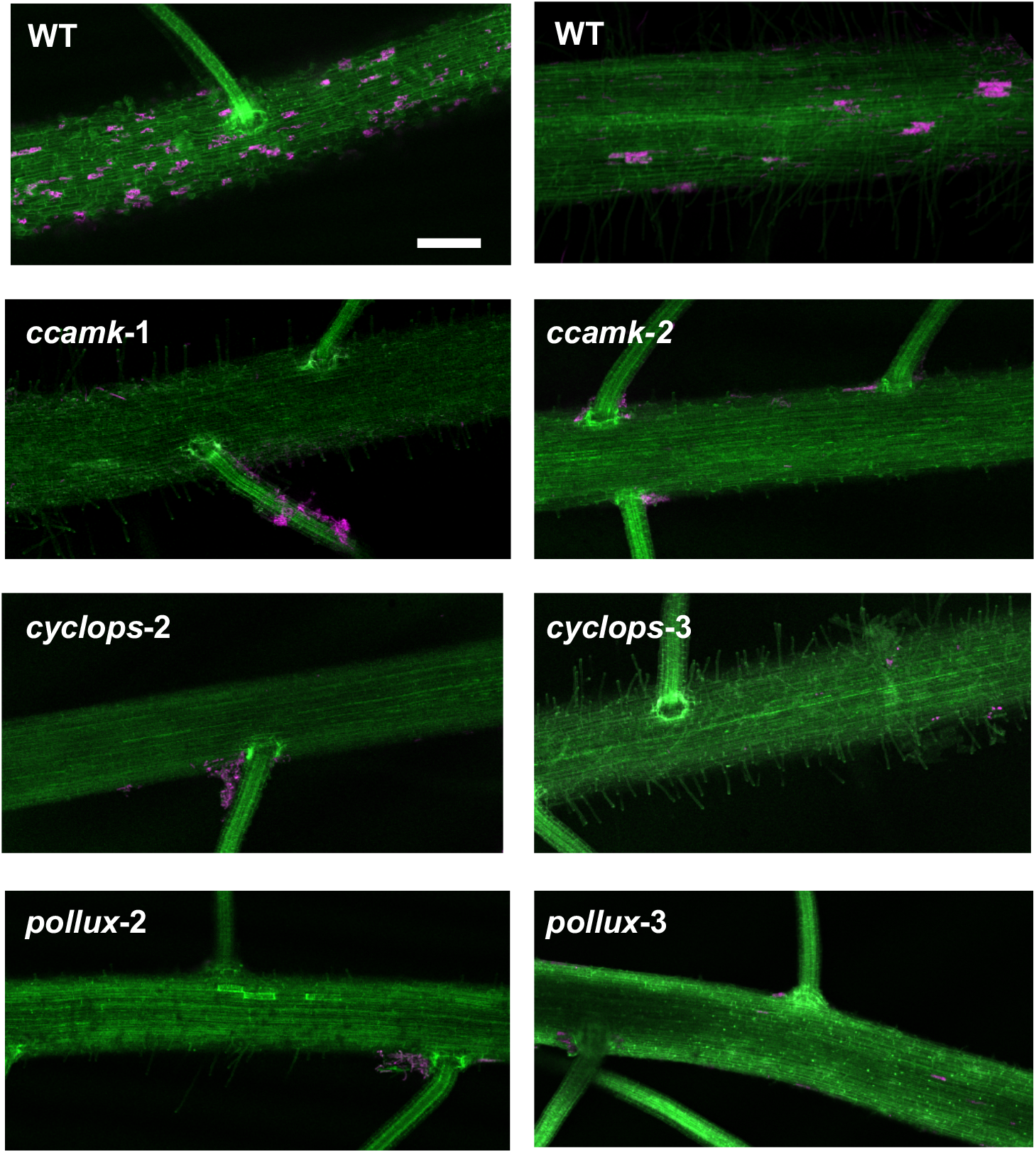
Symbiotic phenotypes of *Oryza sym* mutants. Rice roots inoculated with *Nostoc* at 35 days post inoculation (dpi) were visualized by a confocal microscope. Stacks generated from 90 to 130 frames are shown. In green, suberin autofluorescence from the plant cell walls; in magenta, cyanobacterial chlorophyll autofluorescence. Scale bar: 200 μm.

## 4. DISCUSSION

In this paper, we present data from the first proteomic analysis of the symbiotic interaction between *Oryza* and *Nostoc*. We have identified and quantified protein changes in both organisms at the same time through a novel analytical pipeline, thus facilitating the study of the *Nostoc-Oryza* symbiosis in the host microenvironment. From the whole proteome of both organisms, we have generated species-specific libraries that might be used as reference libraries to future studies. This strategy overcomes traditional proteomic studies in which proteins of both organisms are extracted and analysed separately.

Previous studies of colonization of *Oryza* with *Nostoc* strains showed an early phase of contact that started a few hours of co-culture followed by a second phase of colonization that lasted 20 days (Nilsson et al., 2002; Álvarez et al 2020). This first phase of contact at early stages is critical for the success of the symbiotic process, since is the period when both organisms exchange signals and react to the presence of the partner. We have a total of 1748 proteins differentially expressed along 7days when both organisms remain in contact. This high number of proteins provides evidences of the complexity of the pre-symbiotic crosstalk between *Nostoc* and *Oryza* and importance for the fruitful of the symbiotic process.

This work is the first to show the *Oryza* proteomic changes in response the symbiotic interaction with a cyanobacterium. However, we find similarities with other symbiotic interactions. As in the case of *Oryza*-AM symbiosis, the plant activates expression of three FLAs (Gutjahr et al., 2015). FLAs are cell wall structural glycoproteins that play a crucial role in plant development, cell adhesion and signalling (Seifert, 2018; Wu et al., 2020; Ma and Zhao, 2010). Induction of FLA proteins in response to *Nostoc* might positively influencing accommodation of the cyanobacterium and its interaction with the plant at early stages of the symbiotic process.

Traditionally, plant symbiotic and pathogenic microorganisms have been considered to operate with different modes of action. However, they both induce in the plant signalling components which facilitate colonization and contribute to both symbiotic and pathogenic relationships (Zeilinger et al., 2015; Hentschel et al., 2020). We find overexpressed in *Oryza* three type III peroxidases that might cope with the production of hydroxyl radicals in the plant roots (Table 1). Type III peroxidases have been found induced in rice-AM symbiosis (Guilmil et al, 2005; Gutjahr et al., 2008, 2015). Although their role in the symbiotic process is not yet well understood, they are important for plant cell growth by nonenzymatic loosening of the cell wall during AM infection (Gutjahr et al., 2008). Interaction with beneficial microbes activates in the host plant defence mechanisms in order to facilitate the symbiosis. Thus, glutation S-transferase (GST) enzymes are induced in *Oryza* in response to AM symbiosis to cope with toxic reactive oxygen species (ROS) and lipid hydroperoxides (Gutjahr et al., 2015). Our proteomic analysis revealed the expression in response to *Nostoc* of three GST enzymes. It has been shown recently that *N. punctiforme* induces the expression of GST enzymes in *Arabidopsis* to reduce toxic ROS and limit the extent of the hypersensitive response (Belton et al., 2020). A protein with a Gnk2-homologous domain was also significantly induced in *Oryza* in response to *Nostoc*. Gnk2 domain, similar to calcium-dependent protein kinases from *Arabidopsis*, is found in protein induced by pathogen infection and treatment with ROS or salicylic acid and are involved in the hypersensitive reaction, which is a typical system of programmed cell death (Miyakawa et al., 2009).

In *Nostoc*, 541 proteins were differentially expressed in the presence of *Oryza*. It is important to note that a number of them have been previously described in other *Nostoc*-plant symbioses, reinforcing the robustness of our study. Hence, we find 45 proteins previously highlighted in a supervised machine learning approach of *Nostoc*-Moss symbiosis (Warshan et al., 2017), 8 proteins identified in a previous transcriptomic analysis in response to *Anthoceros punctatus* (Campbell et al., 2008) and 9 proteins in a proteomic analysis of freshly isolated *Nostoc* from the symbiotic gland tissue of the angiosperm *Gunnera* (Ekman et al., 2006). They are involved in signal reception, adhesion, plant cell wall degradation and response to oxidative stress (some of them highlighted in Table 1). The presence of such high number of coincident proteins in the three different *Nostoc*-plant symbiosis might be indicative of the low host selectivity of this symbiotic cyanobacterium. *N. punctiforme*, isolated more than 30 years ago from the cycad *Macrozamia*, is able to stablish both epiphytic and endophytic symbiosis with all four of the major phylogenetic divisions of terrestrial plants (Meeks and Elhai, 2002; Adams et al., 2012). In this line, it is easy to imagine that cyanobacteria, unlike rhizobia, have evolved to sense signals from multiple partners spanning nearly the entire plant kingdom.

Whilst confirming the importance of several characterized proteins and molecular pathways in other *Nostoc*-plant symbioses, this study provides novel evidence to support several new proteins and pathways involved in the pre-symbiotic signalling process. We have found a number of putative Nod-like proteins highly expressed in *Nostoc* in response to the plant, luckily producing lipochitooligosaccharides (LCOs), also known as Nod factors (NFs). They comprise a putative NodB polysaccharide deacetylase, four putative NodE beta-ketoacyl synthases and six proteins with homology to NodO. Reinforcing our results, it has to be noted that *Nostoc* genes with homology to *nodE* had been already detected by Southern blot in previous studies (Rasmussen et al., 1996). *In silico* studies in *Nostoc* have recently identified two ORFs encoding putative homologs of *Rhizobium* NodB and NodC proteins (Gunawardana, 2019). Rhizobial NFs are induced by flavonoids derived from prospective plant hosts (Mus et al., 2016). That might be the case in the *Nostoc*-plant endosymbioses. Induction in *Oryza* of a Flavonoid 3’-monooxygenase involved in the eriodyctiol biosynthesis from naringenin provides first clues of the same regulation in our system. Whether nod genes from *Nostoc* are regulated by plant flavonoids and the role of cyanobacterial Nod factors in the *Nostoc*-plant symbiosis will be addressed in future studies.

NFs play a major role in plant colonization and symbiosis (Mbengue et al., 2020). They produce modifications of the root hair growth leading the formation of pre-infection threads, providing the entrance to the plant. However, this might not be the mode of entry of *Nostoc* into the plant roots. We have observed that the infection started in the plant body and then extended through the root hairs, being root tips unaffected (Álvarez et al., 2020). In legume-rhizobia symbiosis NFs induce at the cellular level depolarization of the cytoplasmic membrane and calcium spiking response, activating genes of the SYM pathway (Kosuta et al., 2008). Genes involved in the SYM pathway are invariantly conserved in all land plant species possessing intracellular symbionts, implying a recruitment of this signalling pathway, independently of the nature of the intracellular symbiont (Radhakrishnan et al., 2020). Our results provide compelling evidence of the requirement of the SYM pathway for *Nostoc* accommodation into the plant roots. However, further studies need to be conducted to understand the role of this pathway during endophytic interactions between *Nostoc* and their host plants.

Overall, this study provides an excellent resource to study proteomic changes in interacting partners without altering the host microenvironment. Our findings reveal molecular pathways activated at earlier stages of the *Nostoc*-rice symbiosis, providing information about the signals produced by these beneficial bacteria which can have long-term implications towards sustainably improving agriculture.

## Acknowledgements

We thank Prof. Uta Paszkowski and Chai Hao Chiu for providing rice mutant lines and for stimulating discussions. Financial support by Fundación General CSIC (program ComFuturo) is also acknowledged.

## Contributions

CA and VM designed and performed the experiments. CA, MB and VM curated the data and interpreted the results. CA and VM conceived the project and wrote the manuscript. FPM-H and MB discussed the results and contributed to the writing. All authors read and approved the final manuscript.

