## Supplementary figures and images for "Early events of the endophytic symbiotic between *Oryza sativa* and *Nostoc punctiforme* involve the SYM pathway"

### Supplemental Figure 1

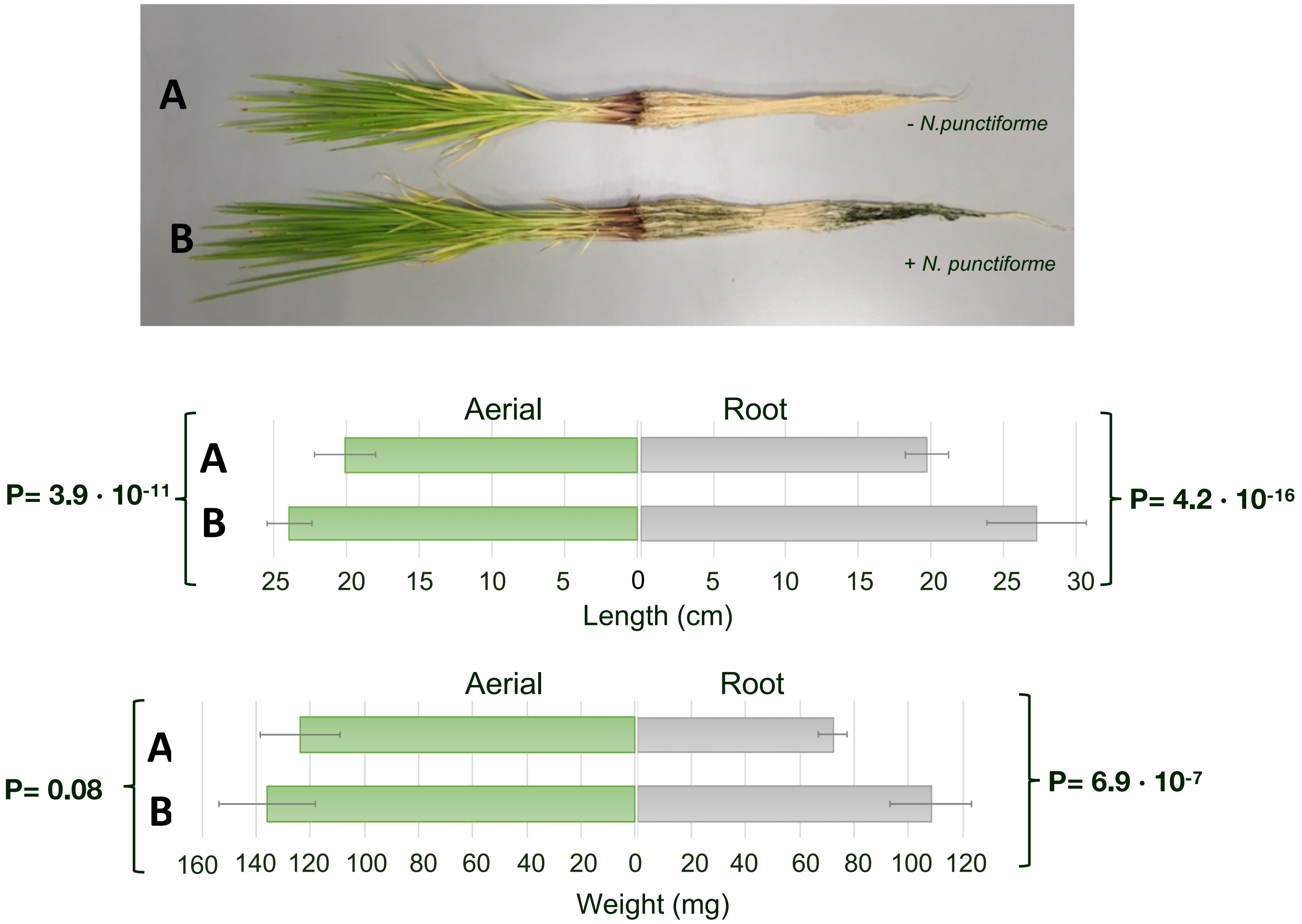


Álvarez et al. Figure S1

### Supplemental Figure 2

*Nostoc*

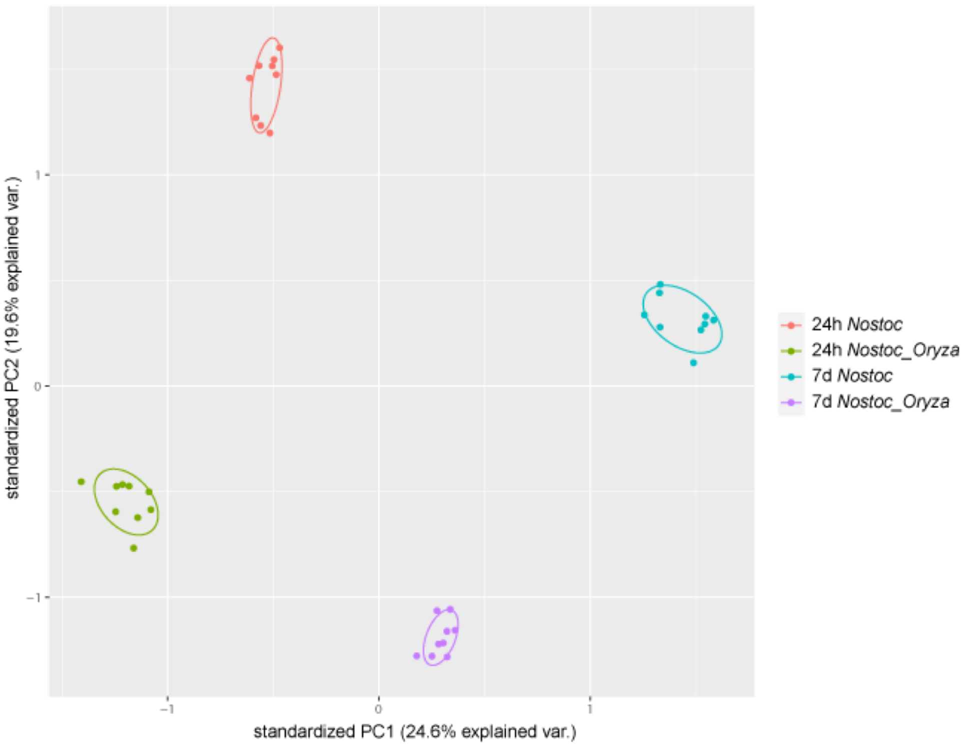


*Oryza*

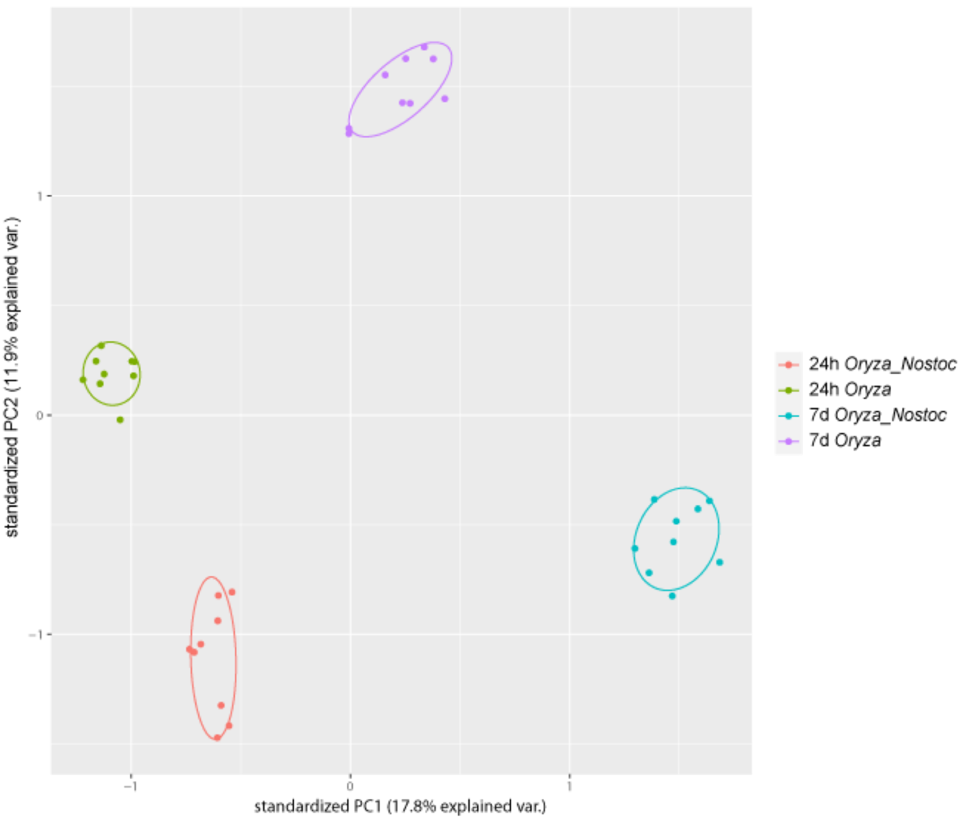


Álvarez et al. Figure S2

### Supplemental Figure 3

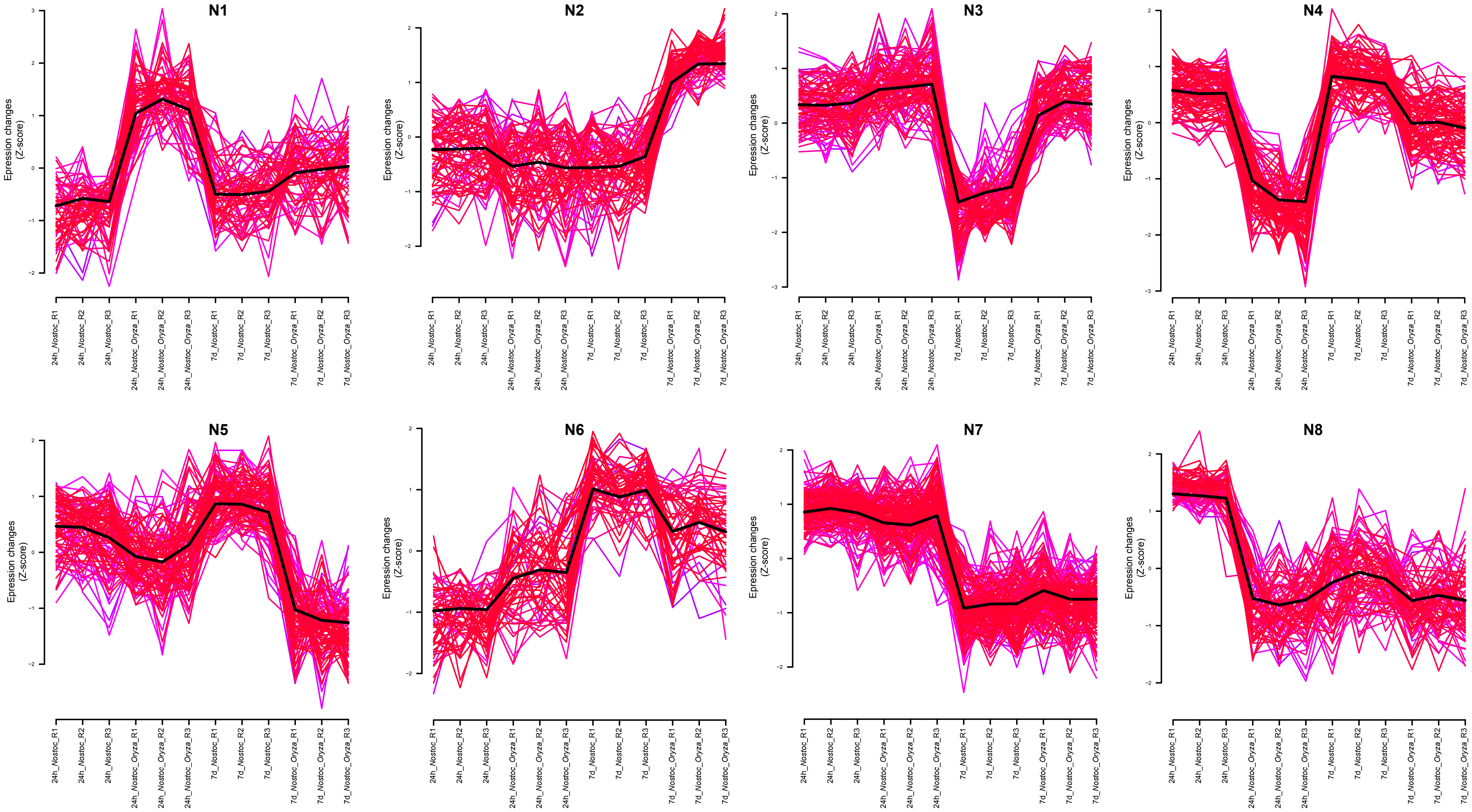

### Supplemental Figure 4

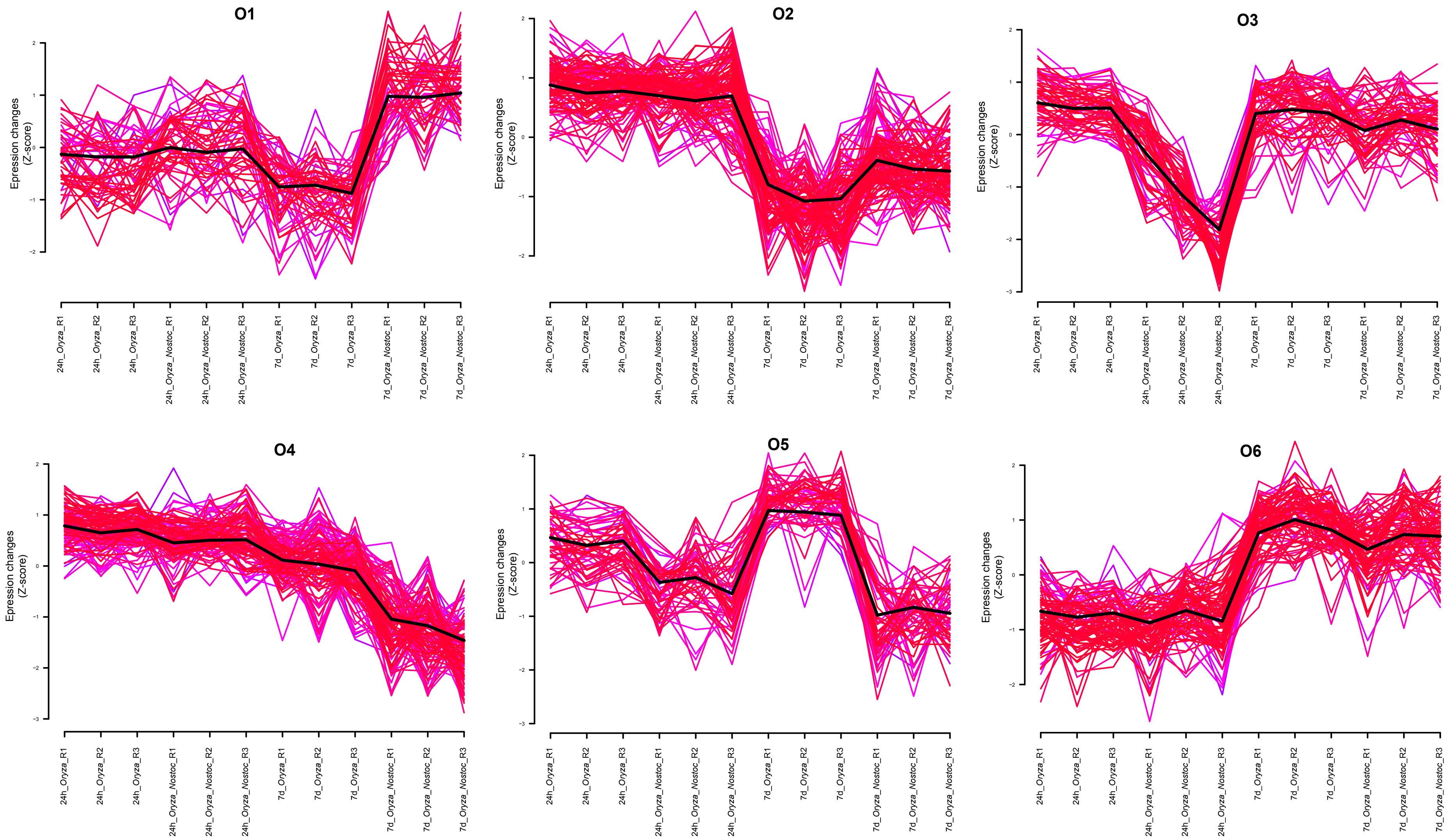
